# Differences in Oligomerization of the SARS-CoV-2 Envelope Protein, Poliovirus VP4, and HIV Vpu

**DOI:** 10.1101/2023.08.18.553902

**Authors:** Julia A. Townsend, Oluwaseun Fapohunda, Zhihan Wang, Hieu Pham, Michael T. Taylor, Brian Kloss, Sang Ho Park, Stanley Opella, Craig A. Aspinwall, Michael T. Marty

**Affiliations:** Department of Chemistry and Biochemistry, University of Arizona, Tucson, AZ 85721, USA; New York Consortium on Membrane Protein Structure, New York Structural Biology Center, New York, NY 10027, USA; Department of Chemistry and Biochemistry, University of California, San Diego, La Jolla, CA 92093, USA; Bio5 Institute, The University of Arizona, Tucson, Arizona 85721, United States

**Keywords:** Viroporins, native mass spectrometry, oligomerization, SARS-CoV-2 Envelope protein, polio VP4, HIV Vpu

## Abstract

Viroporins constitute a class of viral membrane proteins with diverse roles in the viral life cycle. They can self-assemble and form pores within the bilayer that transport substrates, such as ions and genetic material, that are critical to the viral infection cycle. However, there is little known about the oligomeric state of most viroporins. Here, we use native mass spectrometry (MS) in detergent micelles to uncover the patterns of oligomerization of the full-length SARS-CoV-2 envelope (E) protein, poliovirus VP4, and HIV Vpu. Our data suggest that the E protein is a specific dimer, VP4 is exclusively monomeric, and Vpu assembles into a polydisperse mixture of oligomers under these conditions. Overall, these results revealed the diversity in the oligomerization of viroporins, which has implications for mechanisms of their biological functions as well as their potential as therapeutic targets.

## INTRODUCTION

Viroporins are a class of viral proteins that have a wide range of functions in viral life cycles. Although there is very little sequence homology among viroporins from different viruses, several have demonstrated ion channel activity, potentially by forming homo-oligomeric pores in the membrane.^1-3 4^ They also can play vital roles in viral entry and uncoating, in inducing membrane curvature, and in the scission of the viral bud.^5, 6^ Importantly, blocking the channel activity with drugs or deleting viroporins from the viral genome can attenuate viral growth.^4, 7, 8^

The diverse roles of viroporins in infections makes them a potential target for the development of antiviral therapeutics.^9^ Currently, the only viroporin targeted by clinically approved drugs is M2 from Influenza A.^10, 11^ There have been a number of challenges in developing inhibitors for other viroporin targets,^12-14^ including the lack of high-resolution structural information on most viroporins. Viroporins are typically small proteins, with less than 120 amino acid residues.^4^ Most have one or two hydrophobic domains that can form transmembrane alpha-helices within the lipid bilayer.^1, 15^ These hydrophobic domains are thought to drive oligomerization essential for formation of pores within membranes.^16^ However, the oligomeric state of many viroporins has not yet been defined. Without knowledge of the architecture of the pore, it can be challenging to design specific inhibitors to block these channels. Thus, although many viroporins are clinically relevant, they remain a relatively unexplored area for therapeutics.^12^

There are several key analytical challenges associated with studying viroporin structure and interactions, including their small size, intrinsically disordered regions, and potential sensitivity to their local hydrophobic environment. Native mass spectrometry (MS) is a powerful tool to study oligomerization of proteins that addresses many of these challenges. Native MS uses nano-electrospray ionization (nESI) with non-denaturing solvent conditions to preserve noncovalent interactions and can keep proteins folded inside the mass spectrometer.^17^ Because each oligomeric species has a unique mass, native MS provides valuable information on polydisperse oligomers.^18-20 21, 22^ We previously leveraged native MS to reveal the unexpected polydispersity of M2 from influenza A,^22^ which suggested that other viroporins may also display complex oligomeric behavior.

Here, we apply native MS to examine the oligomerization of a range of viroporins, including the envelope (E) protein from SARS-CoV-2, VP4 from poliovirus, and viral protein U (Vpu) from HIV. The SARS-CoV-2 E protein is known to self-assemble within bilayers and is believed to specifically transport cations, including K^+^, Na^+^, and Ca^2+^.^23, 24^ Previous structural studies of the transmembrane domain of the E protein suggest that it forms a pentamer.^25^ However, more recent reports indicate that the various transmembrane constructs of E protein form a dimer in lipid bilayers.^26^ Here, we use native MS to investigate the complexities of the oligomerization of the full-length E protein in diverse chemical environments.

Poliovirus VP4 self-assembles to transport RNA during the viral infection cycle.^27^ Currently, there is little structural information on VP4, but it is believed that the functional form of VP4 is an oligomer.^28^ Previous experiments using chemical crosslinking and SDS-PAGE suggest that VP4 assembles into a hexameric complex.^29^ Myristoylation of the N-terminus is believed to be important in driving oligomerization and function.^30^ However, there remain many unanswered questions surrounding the oligomerization of VP4 and how it may be influenced by its environment.

Finally, HIV Vpu is known to self-assemble and selectively transport monovalent cations across the bilayer.^31^ Several different oligomeric states of Vpu have been proposed, ranging from dimers to octomers,^32^ although it is most commonly described as a pentamer.^33-36^ However, currently there are no structures of Vpu that have been solved as an oligomeric complex, and very little structural data exists for the full-length Vpu, even in its monomer form. Thus, there is still some debate around the oligomerization of Vpu.^37^

Using native MS, we explored their oligomeric state distributions of the full-length forms of the E protein, VP4, and Vpu in a wide range of chemical environments, including different types of detergents, solution pH, ionic strength, and temperature. A range of oligomeric states of these proteins are observed, ranging from the dynamic and polydisperse stoichiometries of Vpu to the more specific dimeric complexes of the E protein. Together, the results contribute to revealing the diverse patterns of self-assembly of viroporins.

## METHODS

### Protein Expression and Purification

#### E Protein Expression and Purification

As previously described,^38^ the plasmid for the KSI-E protein (UniProt ID: P0DTC4) fusion protein was transformed into C43(DE3) *E. coli* cells and grown in LB media to an optical density (O.D.) of 0.5–0.6. Overexpression was induced with isopropyl ß-D-1-thiogalactopyranoside (IPTG) at a final concentration of 1 mM for 3 h at 37 °C before harvesting cells. Cells were then resuspended in lysis buffer (20 mM HEPES, 500 mM NaCl, pH 7.8). Cells were lysed using a LM20 Microfluidizer High Sheer Homogenizer. After lysis, Triton X-100 was then added to a final concentration of 1% (v/v) to the lysate and the solution was stirred at room temperature for 1 hour. Lysate was then clarified by centrifugation at 48,380×g for 30 min at 4 °C, and the supernatant was discarded. The pellet remaining after centrifugation contains inclusion bodies with the protein of interest. This pellet was then resuspended with solubilization buffer (20 mM HEPES, 500 mM NaCl, 1% Fos-choline-16, 1 mM TCEP, pH 7.8) and stirred at room temperature for at least 2 h, or until the inclusion bodies are completely solubilized. The solubilized pellet was then centrifuged at 40,000×g for 30 min at 15 °C, and the supernatant was collected and filtered.

Prior to purification, a HisTrap HP 5 mL column was equilibrated with 10 column volumes of binding buffer (20 mM HEPES, 500 mM NaCl, 20 mM imidazole, 0.1% Fos-choline-16, pH 7.8). The sample was then loaded to a 5 mL HisTrap HP column (GE Healthcare) and washed with approximately 10–20 column volumes of the binding buffer. Afterwards, the column was washed with another 10–20 column volumes of washing buffer (20 mM HEPES, 500 mM NaCl, 50 mM imidazole, 0.1% Fos-choline-16, pH 7.8). The sample was then eluted from the column with HPC elution buffer (20 mM HEPES, 500 mM NaCl, 500 mM imidazole, 0.1% Fos-choline-16, pH 7.8). Eluted protein was pooled and then dialyzed overnight at room temperature with a 10 kDa membrane for against 4 L of dialysis buffer (20 mM HEPES, 50 mM NaCl, 1 mM EDTA, pH 7.8). The next day, thrombin was added to cleave the fusion protein at a final concentration of 10 U/mg and incubate again overnight.^39^

After cleavage of the fusion protein, a reverse nickel purification was performed with a HisTrap HP 5 mL column to separate the histidine tagged KSI protein from the E protein. The column was equilibrated with the binding buffer, and the sample was loaded to the column with the flowthrough collected. The purity of the sample was confirmed with sodium dodecyl sulfate-polyacrylamide gel electrophoresis (SDS-PAGE) and MS. The final sequence and mass of each protein is provided in **Table S-1**.

#### VP4 Expression and Purification

Poliovirus VP4 (UniProt ID: Q84868) with a TEV-cleavable 10×His-MBP-tag was overexpressed in T7 Express pLysY Competent *E. coli* (High Efficiency). Cells were grown at 37 °C in terrific broth media (Thermo Fisher Scientific) to an O.D. of 0.6–0.8. Overexpression was induced with 1 mM IPTG for 3 h, and cells were harvested by centrifugation. Cells were resuspended in lysis buffer (150 mM NaCl, 50 mM Tris, 20 mM imidazole at pH 7.4) containing protease inhibitor. After resuspension, cells were lysed as described above. We found that the MBP-VP4 construct was soluble, so no detergent was added until after cleavage of the MBP tag (see below). The lysate was clarified by centrifugation at 25,000×g for 20 min at 4 °C. Prior to protein purification, a HisTrap HP 5 ml column was equilibrated with 10 column volumes of loading buffer (150 mM NaCl, 50 mM Tris, 20 mM imidazole, pH 7.4). The sample was then filtered using 0.45 μm PES filter, loaded to the column, and washed with 60 column volumes of buffer A. To remove any nonspecific protein binding, the column was washed with 30 column volumes of 5% elution buffer (150 mM NaCl, 50 mM Tris, 400 mM imidazole, pH 7.4). His-MBP-VP4 was then eluted with 100% elution buffer. It was then concentrated using a 10k MWCO filter at 4,000×g for 30 minutes at 4 °C. Dodecyl-β-D-maltoside (DDM, Anatrace) 0.025%, 5 mM BME, and 2 mg/ml TEV (1:50 TEV to protein) were added to the concentrated His-MBP-VP4, and the mixture was then transferred into a 2k MWCO dialysis cassette for overnight dialysis against 150 mM NaCl, 40 mM Tris, and 20 mM imidazole at pH 7.4.

After cleavage of the fusion protein, a reverse nickel purification was performed with a HisTrap HP 5 mL column to separate the histidine tagged MBP from VP4. The column was equilibrated with loading buffer containing 0.025% DDM. The sample was then loaded to the column and the flowthrough of the cleaved VP4 was collected. Residual salts from the protein purification process were removed using a C18 column with a gradient run from HPLC buffer A (0.1% TFA in water) to HPLC buffer B (0.1% TFA in acetonitrile). The final VP4 product was confirmed by SDS-PAGE and MS. Similar results were obtained by purifying the protein without organic solvents and HPLC, but these spectra showed significant sodium adducts during MS.

### Vpu Expression and Purification

Full-length Vpu (UniProt ID: P05919) was transformed into pLysY *E. coli* cells and grown in terrific broth to an O.D. of 0.6–0.8. Overexpression was induced with IPTG at a final concentration of 1 mM for 2 h at 37 °C. Cells were then harvested by centrifugation. Cell pellets were resuspended in a lysis buffer containing 50 mM Tris, 150 mM NaCl, 20 mM imidazole, 0.1% DDM detergent, as well as protease inhibitor. Resuspended cells were stirred at 4 °C for 3–4 h and then lysed as above. Lysate was then centrifuged at 48,380×g for 20 min. Lysate was clarified using a 0.2 μm pore size filter. Prior to protein purification, a HisTrap HP 5 mL column was equilibrated with loading buffer (50 mM Tris, 150 mM NaCl, 20 mM imidazole, and 0.05% DDM, pH 7.4). Sample was then loaded onto a column and washed with 10–15 column volumes of loading buffer. Nonspecific binding was reduced by washing the column with buffer containing 5% elution buffer (50 mM Tris, 150 mM NaCl, 0.4 M imidazole, and 0.025% DDM, pH 7.4). The sample was then eluted from the column with 100% elution buffer. The 10x histidine tag was then cleaved from Vpu by incubating the eluted protein with TEV protease overnight at 4 °C. A reverse nickel purification was performed the next day to remove the TEV and cleaved His tag. The purity of the sample was confirmed with SDS-PAGE and MS.

### Mass Spectrometry Sample Preparation

A series of ammonium acetate solutions were prepared for native MS. All solutions were prepared at a concentration of 0.2 M ammonium acetate, unless stated otherwise. The pH of solutions was adjusted to 5, 7, and 9 using acetatic acid or ammonium hydroxide. Detergents lauryldimethylamine-N-oxide (LDAO), tetraethylene glycol monooctyl ether (C8E4), dodecylphosphorylcholine (DPC) were purchased from Anatrace. Triton X-100, which was purchased from Sigma Aldrich. All solutions were prepared by adding twice the critical micelle concentration (CMC) of the detergent, unless stated otherwise. Viroporins were exchanged into each of these ammonium acetate solutions using BioSpin 6 columns (Bio-Rad) and diluted to a final protein concentration of 20 μM (per monomer) for the E Protein, Vpu, and VP4. Buffer exchanged protein samples were allowed to equilibriate at room temperature for several min prior to analysis, but no change was observed over time. Additional details on native MS analysis, including instrumental parameters, are included in the Supporting Methods along with experimental methods for the electrophysiology experiments.

## RESULTS AND DISCUSSION

Here, we investigated three viroporins—the E protein from SARS-CoV-2, VP4 from poliovirus, and Vpu from HIV— in a wide range of chemical environments to study the effects of chemical environment on the oligomerization of this class of proteins. The chemical environment was altered by changing the solution conditions of the protein prior to ionization (**Figure 1A**). Native mass spectrometry was performed on each sample (**Figure 1B**), and the spectra was deconvolved (**Figure 1C**) to determine the relative intensities of different oligomeric states (**Figure 1D**). The oligomeric state of the protein was determined by dividing the measured complex mass by the mass of the monomeric unit (**Table S-1**). Due to differences in ionization efficiency^40^ and ion transmission,^41, 42^ the native MS intensities do not necessarily directly mirror the concentration of each oligomer species in solution, but the distrubution in native MS signal intensities provides a qualitative picture of the oligomeric states present. Screening the oligomeric states of viroporins over a range of chemical condition provides insights on the structure and interactions of these viroporins.

**Figure 1.**
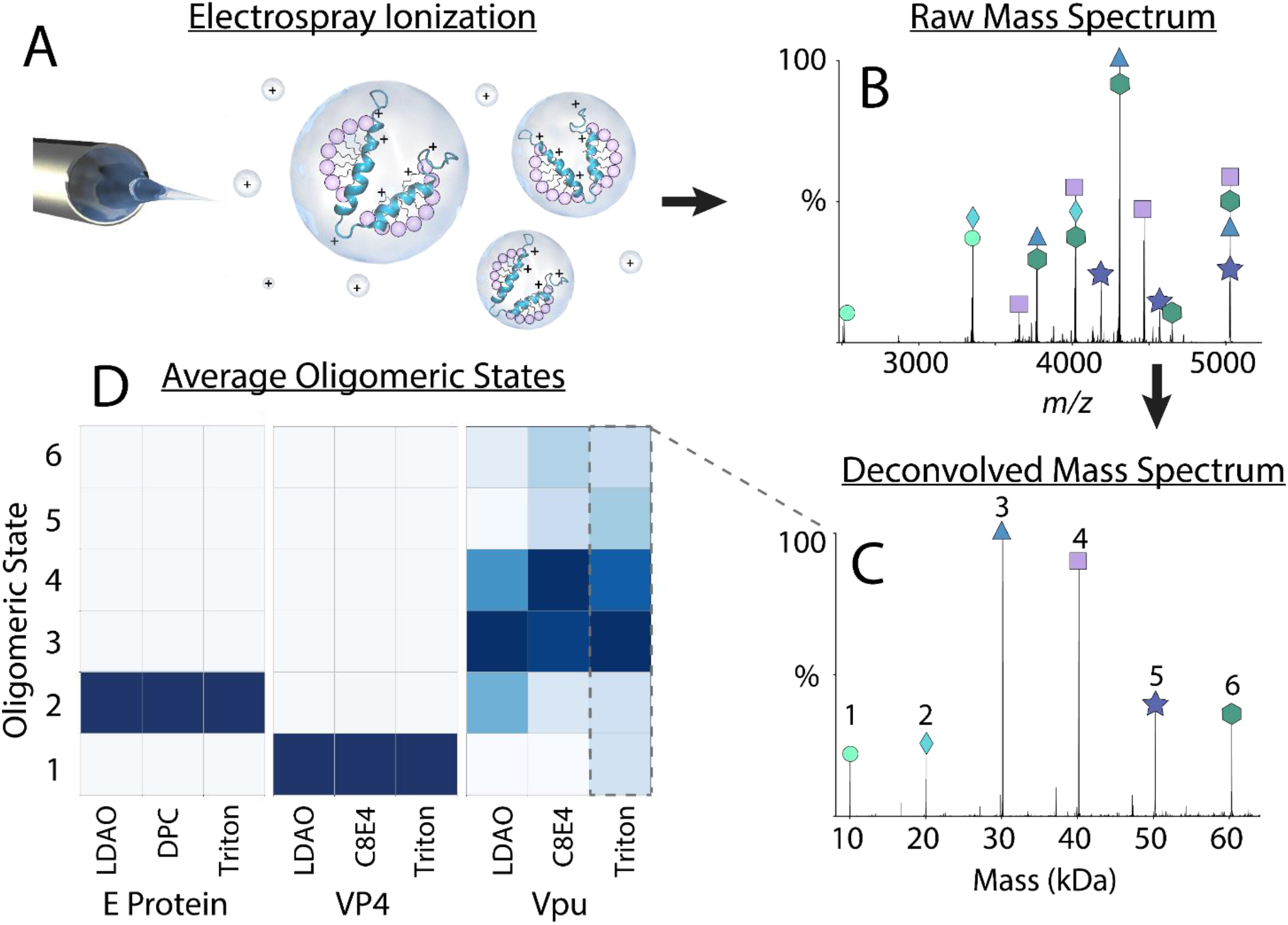
Workflow illustrating the process of studying the oligomeric states of viroporins in various detergents. Electrospray ionization (A) is performed on the protein in detergent, yielding a raw mass spectrum of the sample (B), which is deconvolved into a zero-charge mass spectrum (C). The resulting peak intensities for different oligomers across different proteins and detergent conditions are averaged and displayed on a grid plot (D).

### SARS-CoV-2 E Protein

First, we performed native MS on the E protein in several different detergent conditions. For these initial experiments, we added twice the CMC of each detergent. We discovered that the E protein assembled exclusively as a dimer in LDAO, DPC, and Triton X-100 detergents (**Figure 1D**). Under each of these detergent conditions, we observed residual Fos-Choline-16 detergents from the initial protein purification bound to the E protein. Together, native MS revealed that the E protein forms specific dimers, which was in agreement with the previous structural studies from Zhang, *et al*.,^26^ but did not align with the pentamer previously found by Somberg, *et al*.^25^

Next, we tested the influence of solution pH on the oligomeric state of the E protein. We analyzed the E protein at pH values of 5.0, 7.0, and 9.0. The pH of the solution did not influence E protein oligomerization (**Figure S-1**), and only dimer was observed in all conditions. Because the E protein has three cysteine residues, we also compared the E protein in both reducing and non-reducing conditions at neutral pH. Under both reducing and non-reducing conditions, the E protein behaved as a dimer.

We also investigated the influence of ionic strength on oligomerization. We did not measure any difference in the oligomeric state in the E protein under lower (0.05 M ammonium acetate) or higher (up to 1 M ammonium acetate) ionic strength conditions. However, the number of adducted Foscholine-16 detergents on the protein diminished under higher ionic strength conditions (**Figure 2**). This higher concentration of ions likely outcompetes the adducted Foscholine-16.

**Figure 2.**
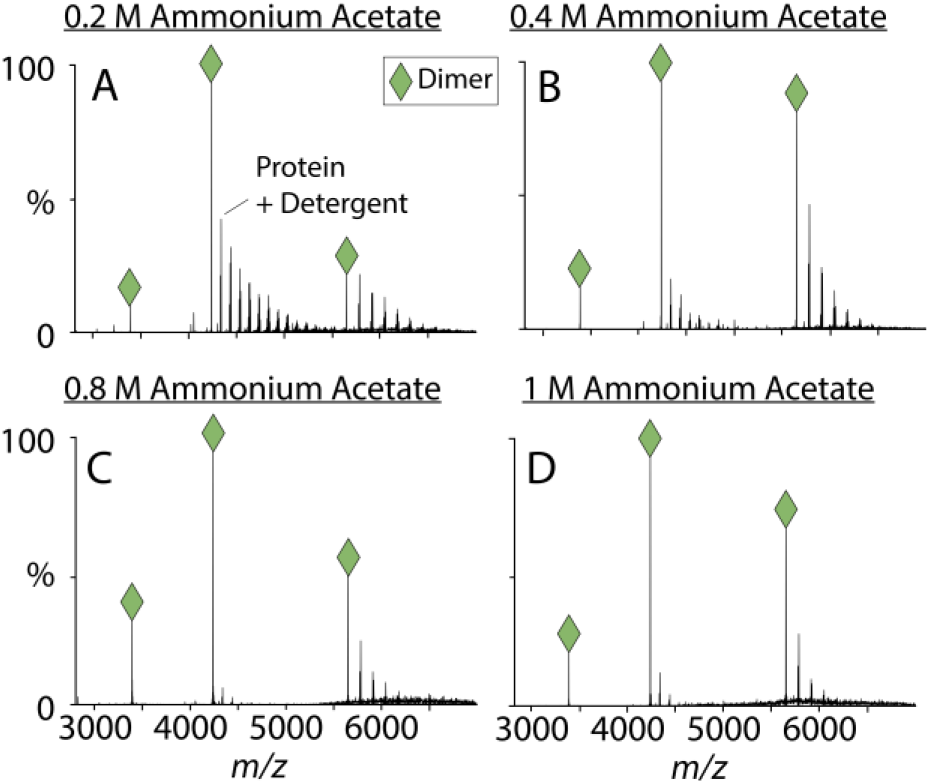
Representative raw mass spectra of the E protein solubilized in LDAO detergent (0.05%) at the concentration of ammonium acetate varied in each sample (0.2 ‒ 1 M ammonium acetate). Charge states for the dimer are marked with a green diamond. Other peaks are from residual detergent bound to the dimer.

Next, we investigated the effect of detergent concentration on the oligomerization of the E protein by adjusting the concentration of LDAO from 0.5 ‒2×CMC. At lower detergent concentrations, we discovered a small peak that corresponded to the mass for the E protein as a trimer, as shown in **Figure S-2**. This E protein trimer was not measured under any other conditions tested and has not been previously reported to our knowledge. Thus, lower concentrations of detergent can affect oligomerization of the E protein, which may indicate the lower levels of lipid/detergent per protein could affect its oligomeric state in bilayers, as previously suggested.^26^

To test the stability of this E protein dimer, we increased the solution temperature on the sample with a variable temperature source.^43^ The temperatures applied ranged from 15‒55 °C, with 10 °C steps. The E protein dimer was present during these temperature ramps at lower temperatures. However, the peak for the dimer became less abundant relative to small detergent clusters at 45 °C, and the dimer was no longer present at 55 °C and above (**Figure S-3**). Interestingly, a clear signal for the monomer did not appear at these higher temperatures where the dimer was less abundant. Instead, the signal for the E protein disappeared altogether, which suggests that the unstable monomer might aggregate at these temperatures.

To validate the activity of the E protein, we also performed electrophysiology studies. These studies were performed using single channel recording under a controlled holding potential of -70 mV. The E protein exhibited flux activity with both Na^+^ and K^+^ ions (**Figure S-4A** and **B**), but different step magnitudes were observed throughout the measurement. The predominant magnitudes of step changes in current were ∼1, 2, 3, and 7 pA in BLMs with DPhPC. These differences in step magnitudes suggested that the E protein may form channels of different stoichiometries. Additionally, solutions containing Ca^2+^ instead of Na^+^ or K^+^ did not exhibit stochastic ion flux (**Figure S-4C**).

As a control, the same experiment was also performed with Fos-choline-16 without protein added to the bilayer, which is the same buffer that the E protein was solubilized in prior to incorporation into the bilayer. No ion conductance was measured upon adding this detergent to the bilayer at the same concentration (**Figure S-4D**), which supports that the ion conductance observed results by the E protein.

Some of the earliest structural work of the E protein proposed that it assembles into a pentameric structure.^44^ However, more recent reports found that the E protein is a dimer.^26^ Using native MS, we found that the full-length E protein assembled nearly exclusively into dimers under all detergent conditions tested (**Figures 1** and **2**). The only conditions where other oligomers were measured for the E protein were in solutions that contained lower concentrations of detergent below that of the CMC, where low intensities of trimer were measured (**Figure S-2**). Previously reported structures of the E protein pentamer were characterized using NMR with the E protein reconstituted into lipid bilayers. However, the protein-to-lipid ratio in these samples are much higher than physiological ratios,^45^ which has the potential to drive the assembly of oligomers. These higher oligomers could be driven by similar effects as the trimer we observed at low detergent concentrations (**Figure S-2**). It is also possible that a lipid environment is necessary to assemble the E protein into a pentamer, but the recent NMR structure of the E protein dimer in lipid bilayers^26^ indicates that the dimer structure is not likely an artifact of detergents.

Although we detected a monodisperse dimer by native MS, ion conductance experiments performed on E protein suggested the presence of polydisperse oligomers, due to the differences in magnitudes of steps in conductance across the bilayer (**Figure S-4**). The discrepancies between the native MS results and electrophysiology results may indicate that the detergents selected might not capture the variety of oligomeric states that could be present in a natural lipid bilayer environment. However, recent structural data of the E protein embedded in a bilayer by Zhang *et al*. also indicated dimer formation.^26^ If the dimer is the functional form of the E protein, this raises two possible mechanisms: 1) that the E protein is able to transport ions in its dimeric form with potentially multiple conductance states, or 2) that the E protein dimers could be building blocks for more transient larger channels, potentially functioning more as a general disruptor of membranes rather than as a specific ion channel.^46^ The behavior of the E protein is complex and will require further investigation to better understand the relationship between its oligomerization and ion channel function.

### Poliovirus VP4

Next, we used native MS to investigate the oligomeric states of poliovirus VP4 in a range of chemical environments. Parameters including detergent type, detergent concentration, ionic strength, and solution pH were all screened. Across all the conditions screened, VP4 was a monomer, as shown in **Figure 3**. This lack of oligomerization is unlike that observed for any of the other viroporins characterized with native MS, all of which oligomerized in detergents.

**Figure 3.**
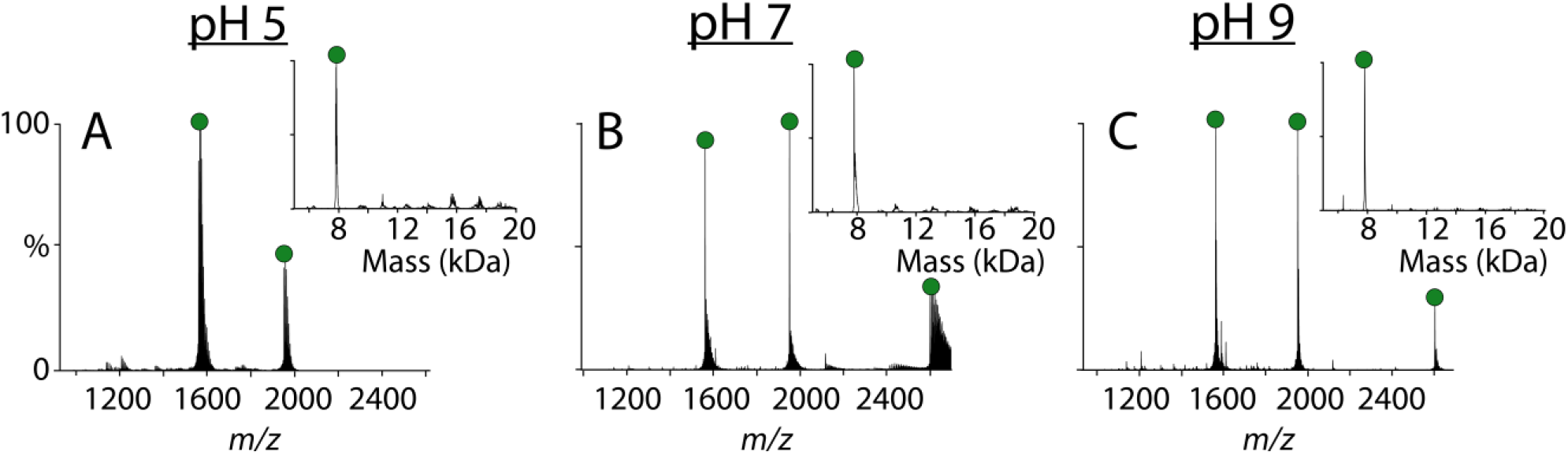
Representative native raw mass spectra of VP4 that has been buffer exchanged into solutions containing 0.2 M ammonium acetate and C8E4 detergent that are pH 5, 7, and 9. The deconvolved mass spectra are inset. Monomer peaks are marked with a green circle.

Prior studies demonstrated that VP4 oligomerizes and transports RNA across the cell membrane.^27^ VP4 is known to be myristoylated on its N-terminal side, and myristylation drives faster assembly of oligomers than unmodified protomers.^47^ Our studies revealed that the unmyristoylated form of VP4 does not form oligomers in these detergent environments. Additionally, VP4 is known to participate in a variety of protein-protein interactions throughout the viral infection cycle, and it is possible that some of these other accessory proteins may be necessary for oligomerization.^48^ Future experiments with the myristoylated proteoform of VP4 will be necessary to better understand these processes, but the current results show no oligomerization of the unmodified VP4 protein in these conditions.

### HIV-Vpu

Finally, we used native MS to study HIV Vpu oligomerization in different chemical environments. We first tested how Vpu was influenced by the type of detergent. When Vpu was solubilized in C8E4 detergent, we observed a mix of oligomers ranging from dimers to hexamers (**Figure 4A**). Similarly, in Triton X-100 detergent, we oligomers ranging from monomers through hexamers are present (**Figure 4B**). However, in LDAO detergent, we observed fewer oligomers, detecting only monomer, dimer, and trimer (**Figure 4C**). These data demonstrate that Vpu oligomerization is sensitive to the detergent environment.

**Figure 4.**
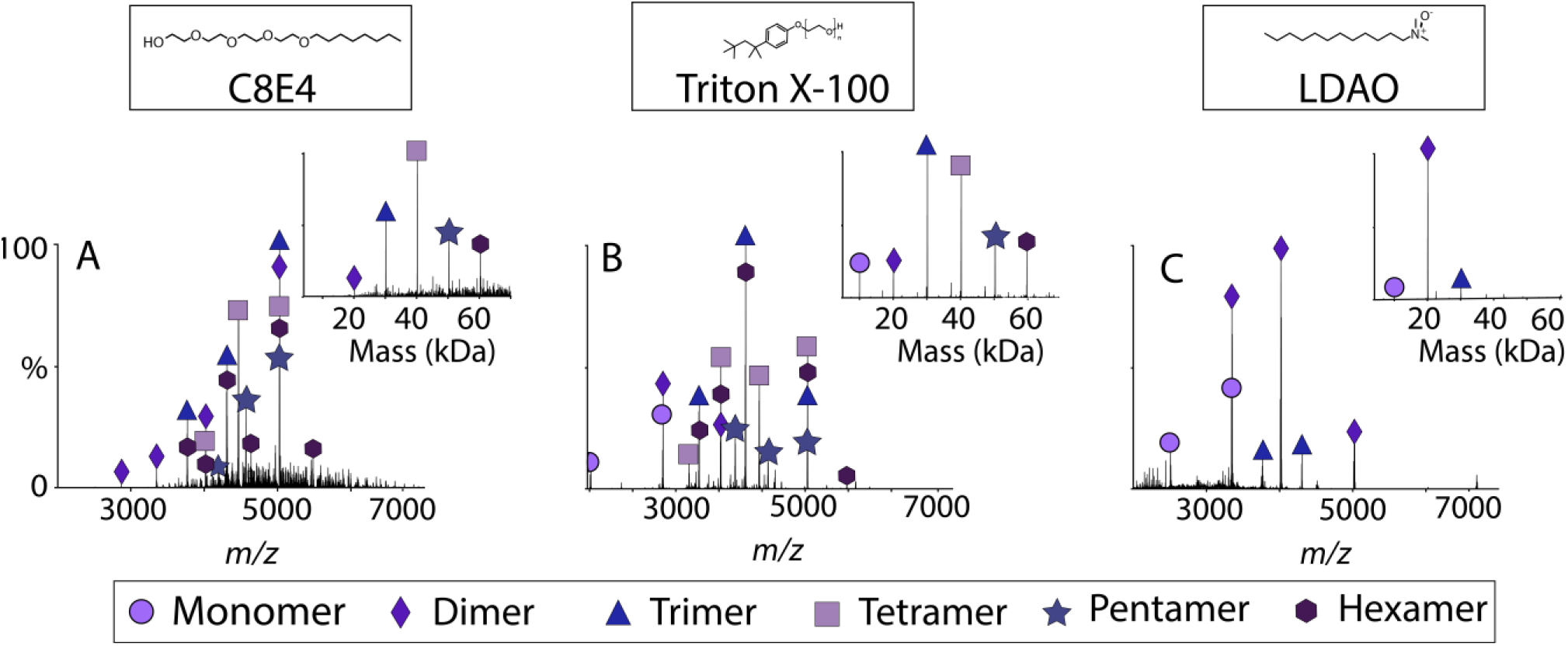
Representative native raw mass spectra of Vpu solubilized in 0.2 M ammonium acetate with (A) C8E4, (B) Triton X100, and (C) LDAO detergents. The deconvolved spectra of each condition are inset, and oligomeric states are annotated with shapes.

Interestingly, although we measured a range of oligomers in all three detergent conditions screened, the oligomeric state profile of Vpu was not significantly influenced by the concentration of the protein. In each of the detergents, we screened the protein in concentrations of 60, 30, and 15 μM, and the oligomeric state profile did not significantly vary across these protein concentrations (**Figure S-5**). We also characterized the oligomeric state of Vpu in a range of solution pH conditions. In LDAO detergent at pH values of 5, 7, and 9, the oligomeric state of Vpu was very similar, as shown in **Figure S-6**. Thus, Vpu oligomerization is not significantly influenced by protein concentration or changes in the solution pH.

To validate the activity of the Vpu while in a bilayer, we performed electrophysiology assays. We first embedded Vpu in BLM’s made up of 1-palmitoyl-2-oleoyl-*sn*-glycero-3phosphocholine (POPC), 1,2-Dioleoyl-*sn*-glycero-3-phosphocholine (DOPC), or 1,2-diphytanoyl-*sn*-glycero-3-phosphocholine (DPhPC) lipids. At fixed membrane potential (V_m_ = -70 mV), the magnitude of current step changes indicates Na^+^ fluxes exhibited a minimum increment of ∼1.5 pA (**Figure S-7A**). The other two predominant magnitudes of the current step changes were ∼2.5 and ∼5 pA, which were consistent among all tested membrane compositions. This result suggests that the Vpu channels exhibit more than one pore stoichiometry. Also, the same predominant step magnitudes indicated that the lipids that were tested had limited effects on the ion permeability of Vpu.

To probe specificity of Vpu ion transport, electrophysiology assays were also performed using buffer containing Ca^2+^ ions instead of Na^+^. Under these conditions, Vpu did not transport Ca^2+^ ions across the bilayer (**Figure S-7B**). Controls were also performed where DDM detergent was added directly to the bilayers instead of Vpu (**Figure S-7C**). No stochastic ion flux events were detected in the absence of Vpu.

Overall, the oligomerization of Vpu appears to be polydisperse and dynamic. A range of oligomers were observed for this protein, and these varied oligomeric states are influenced by the detergent environment. The electrophysiology data of Vpu reveals different step magnitudes for the flux of monovalent cations, which supports the existence of multiple oligomeric states of Vpu. Overall, Vpu exhibited the most complex and polydisperse behavior of the three viroporins investigated here.

Within the literature on Vpu, there is still ongoing debate on its oligomeric state.^31, 49-52^ There is currently no structural information of the full length Vpu forming an oligomer, but there has been one structure solved of full-length Vpu in its monomeric form.^37, 53, 54^ Some previous studies have suggested that Vpu assembles into fixed pentameric complexes,^55^ whereas other studies found that Vpu assembles into a broader range of oligomers.^32, 56-58^ Interestingly, some previous studies suggested that higher solution ionic strengths drive Vpu to assemble into large oligomeric complexes.^56^

Additionally, the electrophysiology assays performed revealed that single channels of Vpu had different magnitudes of ion current across the bilayer (**Figure S-7**). These results suggest that Vpu ion transport behavior is more varied than those of a binary ON/OFF state of canonical ion channels.^59^ Thus, Vpu may form ion channels of different stoichiometries. We hypothesize that the larger channels may exist to enable the transport of larger cations, such as Ca^2+^. However, when the membranes containing Vpu were surrounded by a buffer containing Ca^2+^ instead of Na^+^, no ion conductance was measured. It is possible that these varied oligomers are all specific monovalent cation channels. It could also be that Vpu is sensitive to its local chemical environment and requires a specific lipid environment to form the proper channels, which may have several subtle conformational differences. Further studies will be necessary to better understand the complex physiological roles of the different oligomers of Vpu.

### Comparison of Viroporins

Most of the literature on viroporins suggests that they oligomerize into channels within the lipid bilayer.^1, 60^ Influenza A M2 is the best characterized viroporin, and most of the literature suggests that it forms a specific tetrameric complex that transports protons.^61-64^ Remarkably, native MS performed on M2 in a range of chemical environments, including varied solution pH, different detergents, and a range of different lipid environments revealed a wide variety of previously undetected oligomeric state of M2.^22^ This study raised the question of whether other viroporins from other viruses may also have more varied oligomers and more complexity than initially thought.

Here, we used native MS to study the oligomerization of three different full-length viroporins, the E protein, Vpu, and VP4. We investigated the patterns of oligomerization of these viroporins in different chemical environments, including a range of different detergents, pH conditions, and ionic strength conditions (**Table S-2**). These experiments revealed that the patterns of oligomerization varied widely across proteins, with the E protein forming stable dimers, VP4 remaining monomeric, and Vpu exhibiting highly dynamic behaviors that varied across detergents, as shown in **Figure 5**. Many of these oligomeric states had not previously been detected for these viroporins.

**Figure 5.**
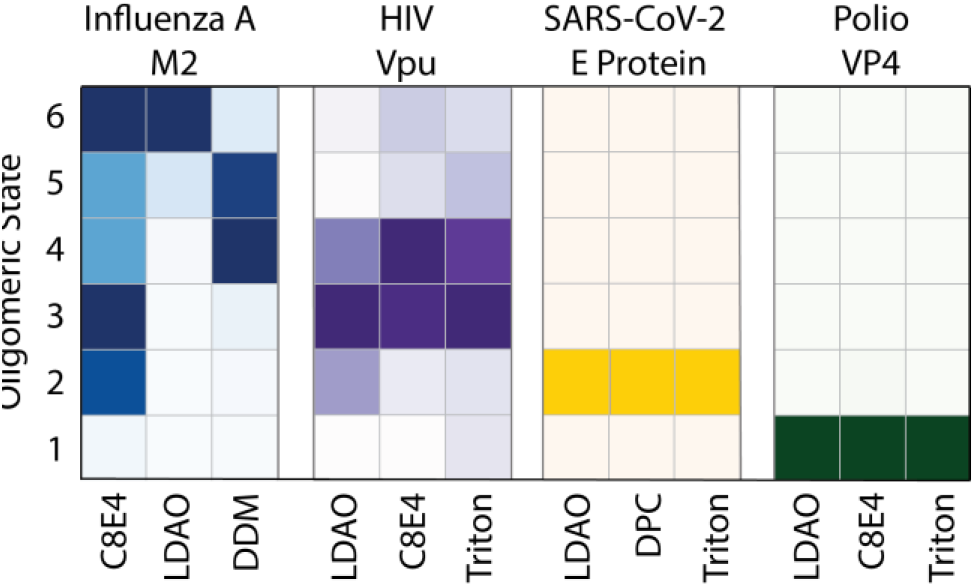
Grid showing the average oligomeric state intensities, ranging from monomer through hexamer, of viroporins M2, Vpu, E Protein, and VP4 in different types of detergent. The darker the color shown, the higher the intensity of the signal for that oligomer.

Our results vary in how they compare with the existing literature on these viroporins. For example, the native MS data on Vpu agrees with some earlier studies. Applying a wide range of analytical tools, Lu *et al*. reported oligomers of various sizes, ranging from monomer through octamer for the transmembrane domain of Vpu.^32^ Our results largely support this polydisperse picture of Vpu oligomerization (**Figures 4, S-5**, and **S-6**).

There is currently much less literature on VP4, so there are no prevailing hypotheses for its oligomerization. However, some earlier experiments with non-polio enteroviruses with analogous forms of VP4 suggest that it oligomerizes into hexameric complexes to transport the genetic material.^65^ However, our results with native MS of polio VP4 revealed only monomers (**Figure 3**). One possible explanation of these differences from the literature is that in our native MS studies, the VP4 used here was not myristoylated.^47, 48, 66^ Future studies using the myristoylated VP4 will be necessary to better understand the oligomerization of VP4, but our results demonstrate the unmodified protein does not oligomerize in these conditions.

The oligomeric state of the E protein remains to be definitively solved. The E protein was initially reported as a pentameric complex,^25^ but a recent NMR structure now suggests a dimeric complex.^26^ Our results with native MS agree with these new results, revealing the assembly of a specific dimer for the E protein **(Figures 2, S-1, S-2**, and **S-3**).

Some of these differences in the literature could be attributed to the challenges with studying the oligomeric states of small membrane proteins, especially for polydisperse mixtures of oligomers.^60^ It is possible that their complexity has been underestimated for several reasons. First, differences in the constructs used, especially between the transmembrane domain versus the full-length protein, may affect oligomeric behavior.^5^ However, our previous results comparing the full-length and transmembrane peptide of M2 revealed similar, although not identical, patterns of oligomerization.^22^

Additionally, differences in sample conditions may significantly affect oligomerization, with changes caused by differences in the buffer, protein concentration, and detergent/lipid environment used. For example, low concentrations of detergent and high concentrations of protein can drive nonspecific oligomers.^67, 68^ However, differences between studies could also indicate true differences in oligomerization caused by different chemical environments, which may be important for physiology, as discussed below. The differences between the E protein dimer observed by native MS and the ion channel activity observed by single channel recording experiments may support the idea that the lipid environment affects oligomerization, although we cannot rule out other effects. Certainly, both M2 and Vpu have shown significant differences in oligomeric state in different chemical environments (**Figure 5**). Differences between the E protein pentamer and dimer have been proposed to have a functional role by forming different complexes in the host cell versus the assembled viron.^26^

In addition to considering the potential effects of sample conditions, it is also important to consider measurement bias during native MS. One potential artifact is that native MS could be artificially generating oligomers. However, this is unlikely because we only measured VP4 in its monomeric form. Another potential artifact is that native MS could potentially disrupt or break apart specific oligomers. This bias is also unlikely because we measured the E protein as a highly specific dimer and M2 as a specific tetramer under certain circumstances.^22^ A final potential artifact could be that native MS was biasing our results, only allowing us to detect certain oligomer species. However, this artifact is also unlikely because we were able to measure Vpu and M2^22^ as a wide range of oligomers. Moreover, agreement of our native MS data with past studies on Vpu polydispersity and recent results on the E protein dimer,^26^ as well as orthogonal measurements made on M2,^22^ indicate that there are unlikely to be large scale biases in oligomeric states observed during native MS.

Ruling out possible measurement artifacts, our current theory is that these viroporins have diverse oligomerization behaviors that are potentially sensitive to the local chemical environment. There are two possible implications of this theory. The first interpretation is that our data may reveal the potential non-physiological biases surrounding the use of different detergents and other buffer conditions in the analysis of viroporins, in which case the physiological oligomeric state may be very different from what we have measured. In this case, this work emphasizes the need to carefully select membrane mimetics and solvent conditions in the analysis of membrane proteins to avoid potential artifacts.^69-71^

However, another possible interpretation of these results is that the dynamic oligomerization of viroporins may be functionally important. Most of the literature surrounding viroporins suggests that they assemble into these fixed complexes and that these monodisperse assemblies are the only physiological form.^4, 16, 25, 33, 72^ However, most viruses genetically encode a relatively small number of proteins. Due to their small genome size,^73^ as well as high mutation rates and frequently changing environmental fitness conditions,^74^ each viral protein may play a variety of roles in the viral life cycle,^75^ which can be accomplished by creating small protein units that can self-assemble into different oligomers that can carry out different biochemical functions.^76, 77^

It is possible that viroporins may assemble into a range of oligomeric states to carry out these different functions, and that different chemical environments may trigger oligomeric rearrangement.

In either case, the native MS experiments reveal previously underexplored differences in the behavior of viroporin complexes in a range of chemical environments. Among the viroporins characterized here, there are few similarities. There is little sequence or structural homology, and patterns in oligomerization vary. It is possible that many of these novel oligomeric states have their own underexplored biochemical function. Overall, these results indicate that viroporins are far more structurally diverse than expected.

## CONCLUSION

In conclusion, we applied native MS to measure the oligomeric states of viroporins E protein, VP4, and Vpu in different chemical environments. The behavior of these proteins varied widely, with the E protein forming dimers, VP4 not oligomerizing at all, and Vpu forming polydisperse and dynamic oligomers. Many of these oligomeric states have not been previously reported. Characterizing the structures of these viroporins will provide a better understanding of virus biochemistry and may support development of therapeutics in the future.

## SUPPORTING INFORMATION

Supplemental methods for native MS and functional studies; supplemental tables with protein sequences and a table of key results; supplemental figures showing effects of different solution conditions and electrophysiology assay data.

### Author Contributions

J.A.T. and M.T.M. designed the research. J.A.T. and M.T.M. wrote the paper with input from the other authors. J.A.T. and O.F. expressed and purified viroporins and collected mass spectrometry data. Z.W. and C.A.A. designed and performed electrophysiology experiments. M.T.T. and H.P. helped with the VP4 purification. B.K. designed plasmids for Vpu and VP4 and performed initial expression screening. S.P. and S.O. provided the E protein plasmid, developed the E protein purification protocol, and provided initial E protein samples.

### Notes

The authors declare no competing financial interest.

## ACKNOWLEDGEMENTS

The authors thank Maria Reinhardt-Szyba, Kyle Fort, and Alexander Makarov at Thermo Fisher Scientific for their support on the UHMR Q-Exactive HF instrument. This work was funded by the National Institute of General Medical Sciences and the National Institutes of Health (T32 GM008804 to J.A.T. and R35 GM128624 to M.T.M.) and the National Science Foundation (CBET-2003297 to C.A.A.). M.T.T. was supported by R35 GM143120 and the University of Arizona. B.K. was supported by P41 GM116799 (PI Wayne A. Henrickson). S.P. and S.O. were supported by R35 GM122501.

