## Supporting Information for "Differences in Oligomerization of the SARS-CoV-2 Envelope Protein, Poliovirus VP4, and HIV Vpu"

### Supplemental Methods

#### Native Mass Spectrometry

All native MS experiments were performed using a Q-Exactive HF Orbitrap (Thermo Scientific) mass spectromter that has ultrahigh mass range modifications (UHMR), as previously described.[1] Nano-electrospray ionization was performed using homemade borrosillicate needles that were pulled with a P-1000 micropipette puller (Sutter Instruments).

Viroporin samples were ionized in positive ion mode with a spray voltage of 1.1–1.5 kV. E Protein and Vpu samples were analyzed with a range of 2,000–10,000 *m/z*. The trapping gas pressure within the mass spectrometer was set to 3. Activation with 50 V of higher-energy collisional dissociation (HCD) energy was applied. To aid in desolvation, 10–50 V of source fragmentation were also applied to each sample. VP4 samples were analyzed with a range of 500–4,000 *m/z* at a resolution setting of 240,000. E protein and Vpu samples were analyzed at a resolution setting of 15,000. Each sample was buffer exchanged and analyzed three times each. Spectra are shown for a single representative replicate.

The native mass spectra for viroporins in detergent were deconvolved using UniDec, as previously described.[1, 2] The settings for deconvolution of viroporins in all detergent conditions included the application of a curved background subtraction of 10, a mass range of 1,000–70,000 Da, a charge range of 1–50, and a FWHM of 1 *m/z.*

#### Functional Studies

##### Material and reagents

$CaCl_2 \cdot 2H_2O$, 2-[4-(2-hydroxyethyl) piperazin-1-yl]-ethanesulfonic acid (HEPES), toluene, and acetone were purchased from Fisher Scientific. NaCl was purchased from EMD Millipore. $HNO_3$ was purchased from Macron. Cholesterol was purchased from Sigma Aldrich. (Tridecafluoro-1,1,2,2-tetrahydrooctyl)-dimethylchlorosilane (PFDCS) was purchased from Gelest, Inc. 1,2-Dioleoyl-*sn*-glycero-3-phosphocholine (DOPC), 1-palmitoyl-2-oleoyl-*sn*-glycero-3-phosphocholine (POPC), and 1,2-diphytanoyl-*sn*-glycero-3-phosphocholine (DPhPC) were purchased from Avanti Polar Lipids. Borosilicate glass capillaries were purchased from World Precision Instruments.

##### Pipet aperture fabrication and black lipid membrane (BLM) formation

The pipet apertures were prepared from 1.5 mm O.D., 1.0 mm I.D. borosilicate glass capillaries. Cleaned capillaries were pulled with a Sutter P-97 puller. To prepare a ~10 μm aperture and a rounded orifice geometry, the tapers of the glass pipets were cut and fire polished with a Narishige MF-900 microforge. Subsequently, the apertures were surface modified with PFDCS, following a previously established gas phase silanization protocol.[3] Silanized pipets were rinsed and dried before use.

DOPC, POPC, and DPhPC lipids dissolved in chloroform were dried under a gentle stream of argon and lyophilized overnight. Immediately before using, the lyophilized lipids were resuspended in *n*-decane. BLMs were formed following a previously established tip-dip technique.[4] Briefly, the tip of the aperture was moved across the interfaces of air, 10 mg/mL resuspended lipids in *n*-decane, and the recording buffers. Formed BLMs across the apertures were kept submerged in the buffer to reconstitute the viroporins.

##### Current and membrane conductance characterization

The recording buffer composition of the recording bath was consistent for current recordings and membrane conductance measurements (0.5 M NaCl and 2.5 mM HEPES, pH 7.0). To test the selectivity on the transported ion, calcium buffer was also prepared but with other compositions controlled (0.5 M $CaCl_2$ and 2.5 mM HEPES, pH 7.0). Single channel recordings (SCRs) were conducted under voltage bias (-70 mV) and were presented after filtered at 500 Hz. Membrane conductance was measured with a previously established protocol. Briefly, the mean current from 21 different membrane potential states was recorded independently and the conductance was calculated from the I-V correlations. All electrophysiology measurements were performed on a HEKA EPC-10 using PatchMaster software.

### Supplemental Tables

**Table S-1:** The amino acid sequences and monomer molecular weight of all three viroporins analyzed in this study.

| Protein | Sequence | Molecular Weight |
|---|---|---|
| **E Protein** | GSMYSFVSEETGTLIVNSVLLFLAFVVFLLVTLAILTALR LCAYCCNIVNVSLVKPTVYVYSRVKNLNSSRVPDLLV | 8,475 Da |
| **VP4** | SNAMGAQVSSQKVGAHENSNRAYGGSTINYTTINY YRDSASNAASKQDFSQDPSKFTEPIKDVLIKTSPMLN | 7,804 Da |
| **Vpu** | MQPIQIAIAALVVAIIIAIVVWSIVIIEYRKILRQRKIDRLIDRLIER AEDSGNESEGEISALVEMGVEMGHHAPWDIDDLAENLYFQ | 10,026 Da |

**Table S-2:** Table summarizing the trends in oligomerization and behavior across each of the three viroporins characterized with native MS.

| | E Protein | VP4 | Vpu |
|---|---|---|---|
| **Average Oligomeric State** | Dimer | Monomer | Varied |
| **Substrate** | $Na^+/K^+$ | RNA | $Na^+/K^+$ |
| **Influenced by Detergent Type?** | X | X | ✓ |
| **Influenced by Detergent Concentration?** | ✓ | X | - |
| **Influenced by Protein Concentration?** | X | X | X |

### Supplemental Figures

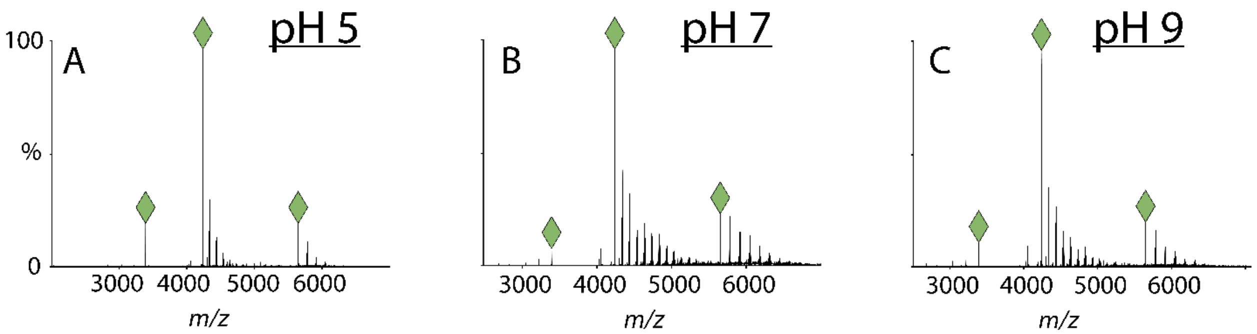

**Figure S-1:** Representative raw mass spectra of the E protein in 0.2 M ammonium acetate solution with pH 5.0 (A), 7.0 (B), and 9.0 (C) and LDAO detergent. Charge states for dimers are marked with green diamonds. The additional peaks in each spectrum correspond to Fos-Choline-16 detergents from the initial purification that remained bound through the buffer exchange.

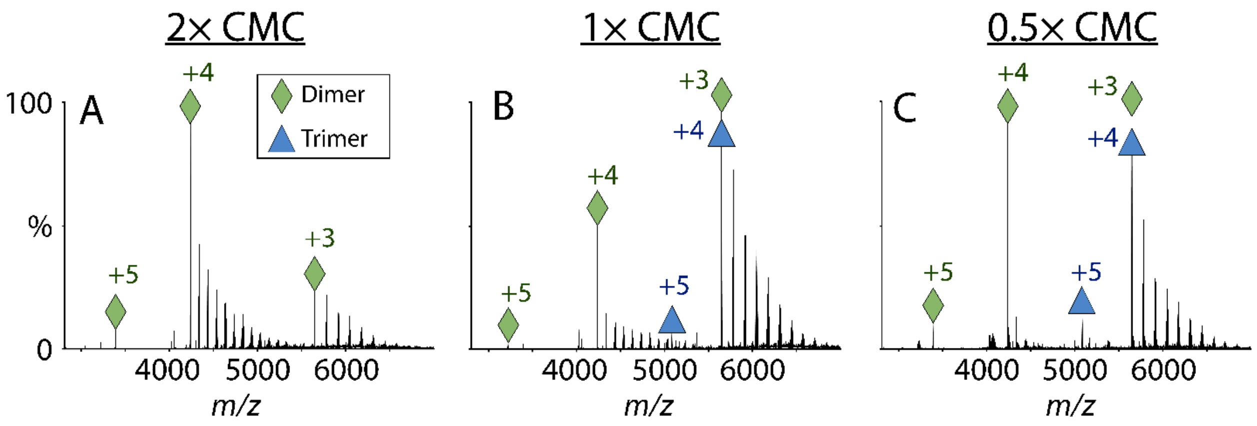

**Figure S-2:** Representative raw mass spectra of the E protein in 0.2 M ammonium acetate with varying concentrations of LDAO detergent, ranging from 2× CMC (A), 1× CMC (B), and 0.5× CMC (C). The CMC of LDAO is 0.025%. Charge states for dimers are marked with green diamonds, and charge states for trimers are marked with blue triangles. Additional detergents bound to the protein in each spectrum are Fos-Choline-16 detergents, residual from the initial protein purification.

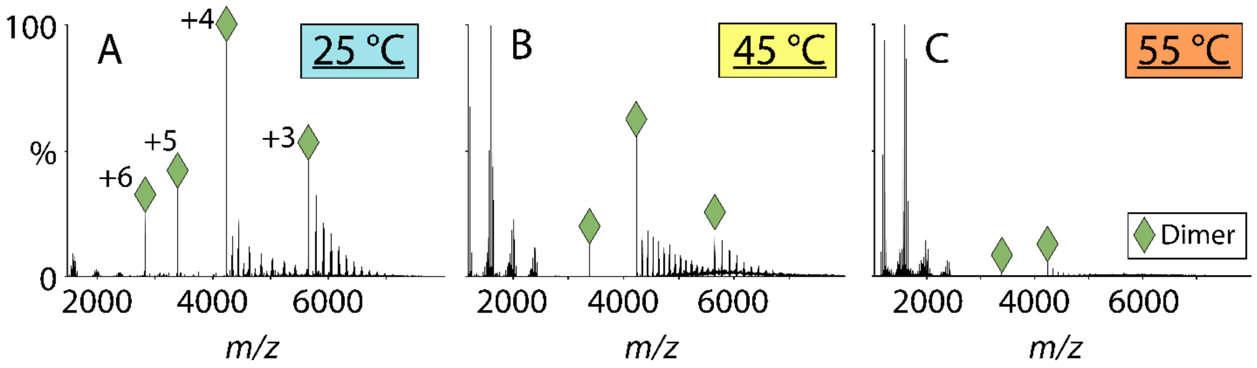

**Figure S-3:** Representative raw mass spectra of the E protein solubilized in 0.2 M ammonium acetate and 0.05% LDAO detergent. The spectra shown show the E protein with the solution temperature at 25, 45, and 55 °C. The charge states of the E protein are labeled under the 25 °C condition. Charge states for dimers are marked with green diamonds. The additional peaks in each spectrum correspond to Fos-Choline-16 detergents from the initial purification that remained bound through the buffer exchange.

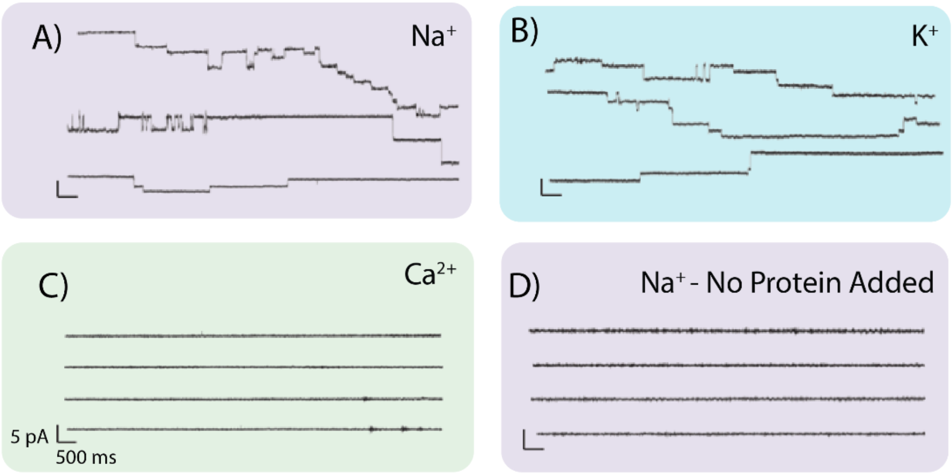

**Figure S-4:** Representative SCR results of E protein in (A) DPhPC BLMs and $Na^+$ buffer, (B) DPhPC BLMs and $K^+$ buffer. Control experiments were performed with (C) $Ca^{2+}$ as the ion in the buffer and with (D) Fos-choline-16 added instead of E protein.

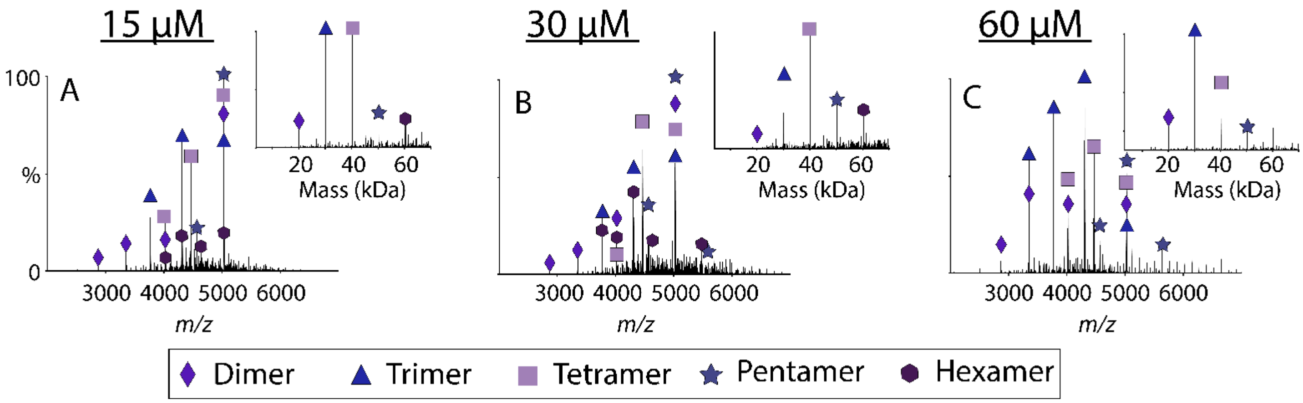

**Figure S-5:** Representative native raw mass spectra of Vpu in 0.2 M ammonium acetate and C8E4 where the concentration of protein monomer is at 15 (A), 30 (B), and 60 µM (C). The deconvolved mass spectra are inset. Different oligomers are marked with different shapes and colors, as shown in the legend.

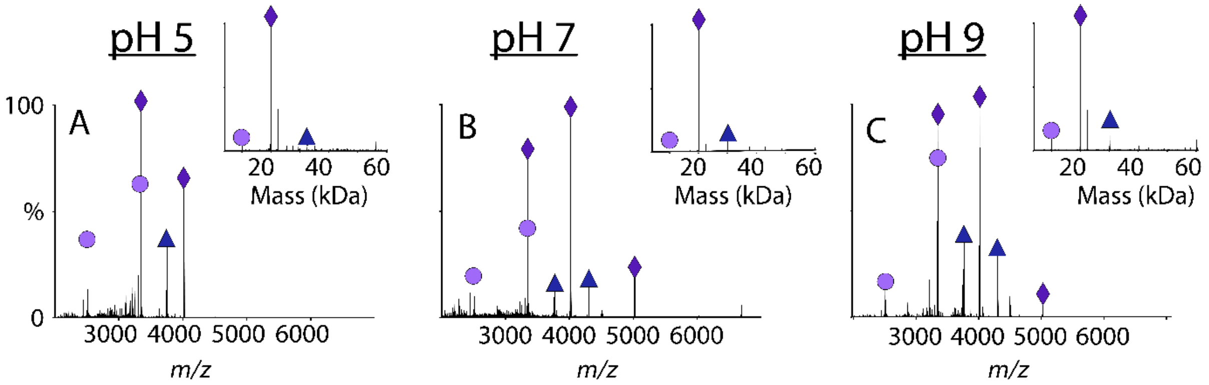

**Figure S-6:** Representative native raw mass spectra of Vpu in 0.2 M ammonium acetate and LDAO where the solution pH is 5.0 (A), 7.0 (B), and 9.0 (C). The deconvolved mass spectra are inset.

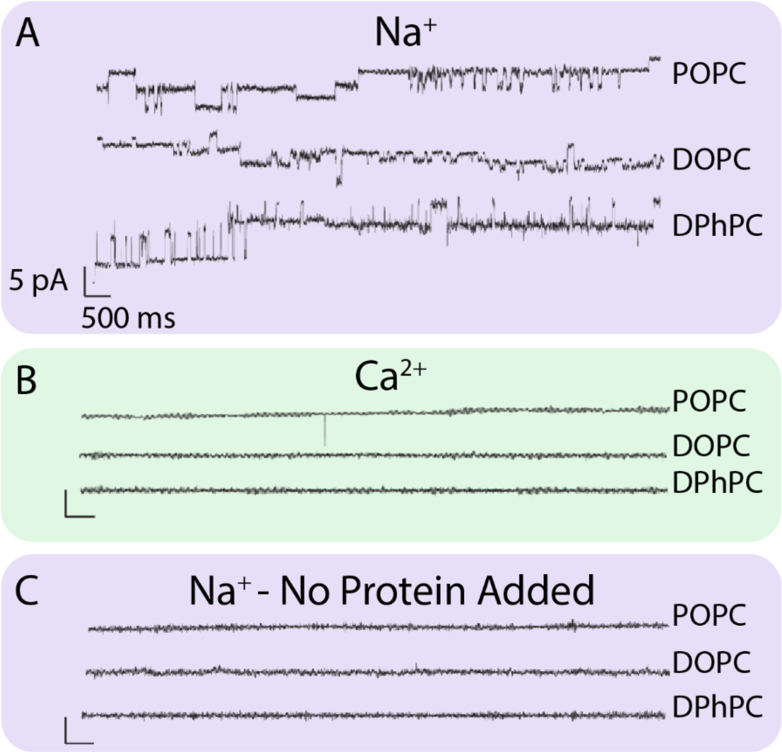

**Figure S-7:** Representative SCR results of Vpu embedded in bilayers made of POPC, DOPC, and DPhPC with A) $Na^{+}$ and B) $Ca^{2+}$ buffer. As a control, C) DDM detergent was added without Vpu to a bilayer with $Na^{+}$ buffer.
